# Co-exposure to Polyethylene Fiber and *Salmonella enterica* Typhimurium Alters Microbiome and Metabolome of *in vitro* Chicken Cecal Mesocosms

**DOI:** 10.1101/2023.11.22.568320

**Authors:** Chamia C. Chatman, Elena G. Olson, Allison J. Freedman, Dana K. Dittoe, Steven C. Ricke, Erica L-W. Majumder

## Abstract

Humans and animals encounter a summation of exposures during their lifetime (the exposome). In recent years, the scope of the exposome has begun to include microplastics. Microplastics (MPs) have increasingly been found in locations where there could be an interaction with *Salmonella enterica* Typhimurium, one of the commonly isolated serovars from processed chicken. In this study, the microbiota response to a 24-hour co-exposure to *Salmonella enterica* Typhimurium and/or low-density polyethylene (PE) microplastics in an *in vitro* broiler cecal model was determined using 16S rRNA amplicon sequencing (Illumina) and untargeted metabolomics. Community sequencing results indicated that PE fiber with and without *S.* Typhimurium yielded a lower *Firmicutes/Bacteroides* ratio compared to other treatment groups, which is associated with poor gut health, and overall had greater changes to the cecal microbial community composition. However, changes in the total metabolome were primarily driven by the presence of *S.* Typhimurium. Additionally, the co-exposure to PE Fiber and *S*. Typhimurium caused greater cecal microbial community and metabolome changes than either exposure alone. Our results indicate that polymer shape is an important factor in effects resulting from exposure. It also demonstrates that microplastic-pathogen interactions cause metabolic alterations to the chicken cecal microbiome in an *in vitro* chicken cecal model.

**IMPORTANCE:** Researching the exposome, a summation of exposure of one’s lifespan, will aid in determining the environmental factors that contribute to disease states. There is an emerging concern that microplastic-pathogen interactions in the gastrointestinal tract of broiler chickens may lead to an increase in *Salmonella* infection across flocks and eventually increased incidence of human salmonellosis cases. In this research article, we elucidated the effects of co-exposure to polyethylene microplastics and *Salmonella enterica* serovar Typhimurium on the ceca microbial community. *Salmonella* presence caused strong shifts in the cecal metabolome but not the microbiome. The inverse was true for polyethylene fiber. Polyethylene powder had almost no effect. The co-exposure had worse effects than either alone. This demonstrates that exposure effects to the gut microbial community are contaminant specific. When combined, the interactions between exposures exacerbate changes to the gut environment. The results herein support current *Salmonella* mitigation efforts and understanding microplastics-pathogen interactions.

## INTRODUCTION

Plastic production has steadily increased in the United States while recycling rates have remained small (1, 2). As a result, non-recycled plastics are disposed of into landfills or as litter in the environment where weathering and biodegradation will break down plastics into smaller plastic fragments called microplastics (3). Microplastics (MPs) are defined as plastic fragments less than 5 mm in diameter (3–5). As MP contamination increases in the environment, the amount of animal and human MP exposure also increases. Interaction with MPs is most likely to occur through food systems, as ingestion and inhalation are common routes of exposure (4, 5). This is evidenced by the detection of MPs in food products like seafood, prepared meats, salt, beer, sugar, and honey (6–8).

Several studies have implicated MPs exposure to reproductive toxicity, increased reactive oxygen species, gut dysbiosis and inflammation in various species. With global plastic pollution to terrestrial environments estimated to be 13 to 25 million metric tons per year (13), terrestrial animals including livestock have increased risk of exposure. However, evaluation of health effects resulting from microplastic exposure must also account for interaction with factors such as microorganisms. In aquatic environments, microbial utilization of MPs or their degradation products as carbon sources has resulted in biogeochemical changes to the system (14). For example, Chen *et al.* (2020) reported that when microorganisms colonized polypropylene MPs there were alterations to the nitrogen and phosphorous cycles in an aquatic system (14,15). MPs-microbe interaction also leads to microbial colonization on MPs and MPs degradation (16). As such, adverse outcomes associated with MP exposure greatly depend on the foreign materials or microorganisms consumed simultaneously and how these factors interact.

Lwanga et. al. (2017) was among the first to demonstrate that plastic pollution could move along the terrestrial food chain and enter the digestive tract of broiler chickens (17). Broiler chickens are the most common breed of meat-producing chicken in the United States (18). Interactions between MPs and pathogens such as *S*. Typhimurium are becoming an emerging concern in poultry production facilities as both contaminants have been detected in processed poultry (6, 7, 19). *Salmonella enterica* serovar Typhimurium (*S.* Typhimurium) is one of the more commonly isolated foodborne pathogens from broiler meat (20, 21). *S.* Typhimurium, a Gram-negative, enteric pathogen, invades intestinal epithelial and is commonly responsible for salmonellosis in human cases (19, 22). Salmonellosis in humans typically occurs after consumption of undercooked meat and results in gastroenteritis, vomiting, and diarrhea (20). Chickens can acquire *S.* Typhimurium from feed, water, air, rodents, or insects (23). In broilers, salmonellosis presents as weakness, ruffled feathers, and weight loss (19, 22). Systemic infection is usually isolated to broilers less than 3 days old and older birds (≥ 3 weeks post-hatch) experience subclinical infection (22, 24). In addition, *Salmonella* infection does not typically present clinical manifestations in chickens unless newly-hatched (24). Overall, the age of the broiler, *Salmonella* serotype, and physiological state of the *Salmonella spp.* influence *Salmonella* persistence in broilers (23).

There are two main routes of transmission of *Salmonella spp.* among flocks. The first is via vertical transmission (i.e., hens to offspring) whereby *Salmonella* invades the reproductive organs and results in infection to the eggs prior to hatch (25). The second is horizontal transmission across flocks due to fecal shedding. Both routes of transmission result in widespread infection and cross-contamination as *S.* Typhimurium can form biofilms (20). Biofilms are defined as cell clusters that are enclosed in a complex matrix of extracellular polymeric substances or EPS (26, 27). The EPS matrix is primarily composed of polysaccharides, extracellular DNA, and proteins which allow fixed bacteria to be protected from environmental stressors within meat processing facilities (26, 28).

Biofilms are an important aspect of food safety as *S.* Typhimurium biofilms are more resistant to antimicrobials commonly used in meat processing facilities (20). Likewise, co-exposure to MPs and this pathogen would provide more surface area and potentially seed *S.* Typhimurium biofilm growth in the gastrointestinal tract of broilers. As a result, there is concern that biofilm formation in the gastrointestinal tract of broilers will result in an increased prevalence of S. Typhimurium in flocks and therefore increased incidence of human salmonellosis cases. Therefore, understanding the interaction of MPs and S. Typhimurium is of the upmost importance.

Despite research indicating the public health risk of MPs-microbe interactions, there are limited studies on the effect of such co-exposure on the gastrointestinal tract. The objective of this study was to elucidate the effects of co-exposure of low-density polyethylene microplastics and *S.* Typhimurium on the chicken cecal microbiome and metabolome. This study focused on low-density polyethylene (PE) because it is one of the most abundant plastics found in the environment and in food packaging materials (8). Low-density PE comes in two common forms: powder and fiber. Using *in vitro* cecal mesocosms for pathogen and MPs co-exposure assessment, it was hypothesized that simultaneous exposure to PE MPs and *S.* Typhimurium would lead to greater disruption in the cecal microbiome and metabolome compared to either contaminant singularly. Effects of PE fiber and PE powder with or without *S*. Typhimurium were first assessed by observing overall taxonomic composition using 16S rRNA gene amplicon sequencing targeting the V4 region. Based on this, the fold change of genera was calculated to quantify changes in relative abundance. Global untargeted metabolomics was conducted and assessed with functional and statistical analyses. The data presented indicates that polyethylene fiber, but not powder leads to altered taxonomic composition, while *Salmonella* dominates small metabolite presence. The results presented here will aid in ameliorating risk assessment standards for livestock and contribute to our general knowledge on MPs-pathogen interactions.

## MATERIALS AND METHODS

### Preparation of Polyethylene Microplastics

Low density polyethylene (LDPE) fibers (GoodFellow, LS554234) were cut to 50 µm using a cryotome (Leica biosystem 1950 cryotome) protocol for preparing microplastic fibers (29). Polyethylene fiber sizes were verified using scanning electron microscopy (Zeiss GeminiSEM 450) (Figure S5). Low density polyethylene powder (Thermo Fisher Scientific, Lot: Z05D030) and polyethylene fiber were weighed (50 mg) and aliquoted into designated glass serum bottles.

### Preparation of Nalidixic acid-resistant *Salmonella*

A frozen, pure culture of *Salmonella enterica* Typhimurium (ATCC 14028) was streaked for isolation on XLT-4 agar (BD Difco; Lot: 1216783) and incubated at 37°C for 24 h. After, one colony was inoculated in 10 mL of Tryptic Soy Broth (TSB) (Bacto; Lot:1131384) and incubated for 24 h at 37°C. A 1:2 nalidixic acid serial dilution was used to produce a nalidixic acid-resistant (NA-resistant) strain of *S.* Typhimurium. Each dilution was inoculated with 1 mL of *S.* Typhimurium and incubated for 24 h at 37°C. Following which the dilution with the highest growth of NA-resistant *S.* Typhimurium was used to continue the serial dilution. This process continued until the *S.* Typhimurium was able to grow in TSB containing 64 µg/mL nalidixic acid. The inoculum was then streaked onto XLT-4 containing 64 µg/mL nalidixic acid (VWR; Lot:21J285302) and incubated overnight at 37°C. Following this one colony was selected and inoculated into 40 mL of TSB and incubated for 24 h at 37°C. The day of the study 200 µL of the NA-resistant *S.* Typhimurium culture was inoculated into designated glass serum bottles.

### Cecal culture preparation and treatment

The *in vitro* cecal microbiome model utilized ceca from 4-week-old Aviagen Ross 308 Broiler chickens from Whelps Hatchery in Bancroft, Iowa. All birds were housed at the University of Wisconsin’s Poultry Research Facility in floor pens and fed commercial grade corn and soybean double pelleted chicken diet prepared onsite at the University of Wisconsin-Madison (Table S7). This study was conducted in accordance with the recommendations from the Guide for the Care and Use of Laboratory Animals. All protocols were reviewed and approved by the University of Wisconsin-Madison’s Institutional Animal Care and Use Committee (IACUC ID: A006627). The 10 broiler chickens selected for this study were humanely euthanized with carbon dioxide asphyxiation. Ceca from each bird were separated at the ileal-cecal junction with alcohol-dipped and flame-sterilized tools and placed in sterile samples bags which were then placed in anaerobic boxes for immediate transport to the anaerobic chamber.

Under anaerobic conditions in a Coy (Coy Laboratory Products, Grass Lake, Michigan, USA) anaerobic chamber with atmosphere containing 5% O_2_, 10% CO_2_, 85% N_2_, 0.1 g of cecal content from the proximal end of each cecum was collected into sterile 2 mL centrifuge tubes and resuspended in 900 µL of anaerobic dilution solution (0.45 g/L K_2_HPO_4_, 0.45 g/L KH_2_PO_4_, 0.45 g/L (NH_4_)2SO_4_, 0.9 g/L NaCl, 0.1875 g/L MgSO_4_-7H2O, 0.12 g/L CaCl_2_-2H_2_O, 0.0001% resazurin, 0.05% cysteine-HCl, and 0.2% NaCO_3_) (30, 31) (Figure 1). The resuspended cecal samples from each bird were then diluted 1:3,000 in ADS and aliquoted in 20 mL portions 6 times into serum bottles (6 treatments). In addition, 0.2 g of the chicken feed was added to the digesta diluent as a nutrient source. Serum bottles were then capped and crimped to maintain anaerobic conditions and placed in a stationary incubator at 37°C. The digesta diluent containing poultry basal diet were incubated at 37°C for a 24 h acclimation period prior to treatment inoculation (30–32) (Table S1).

**Figure 1.**
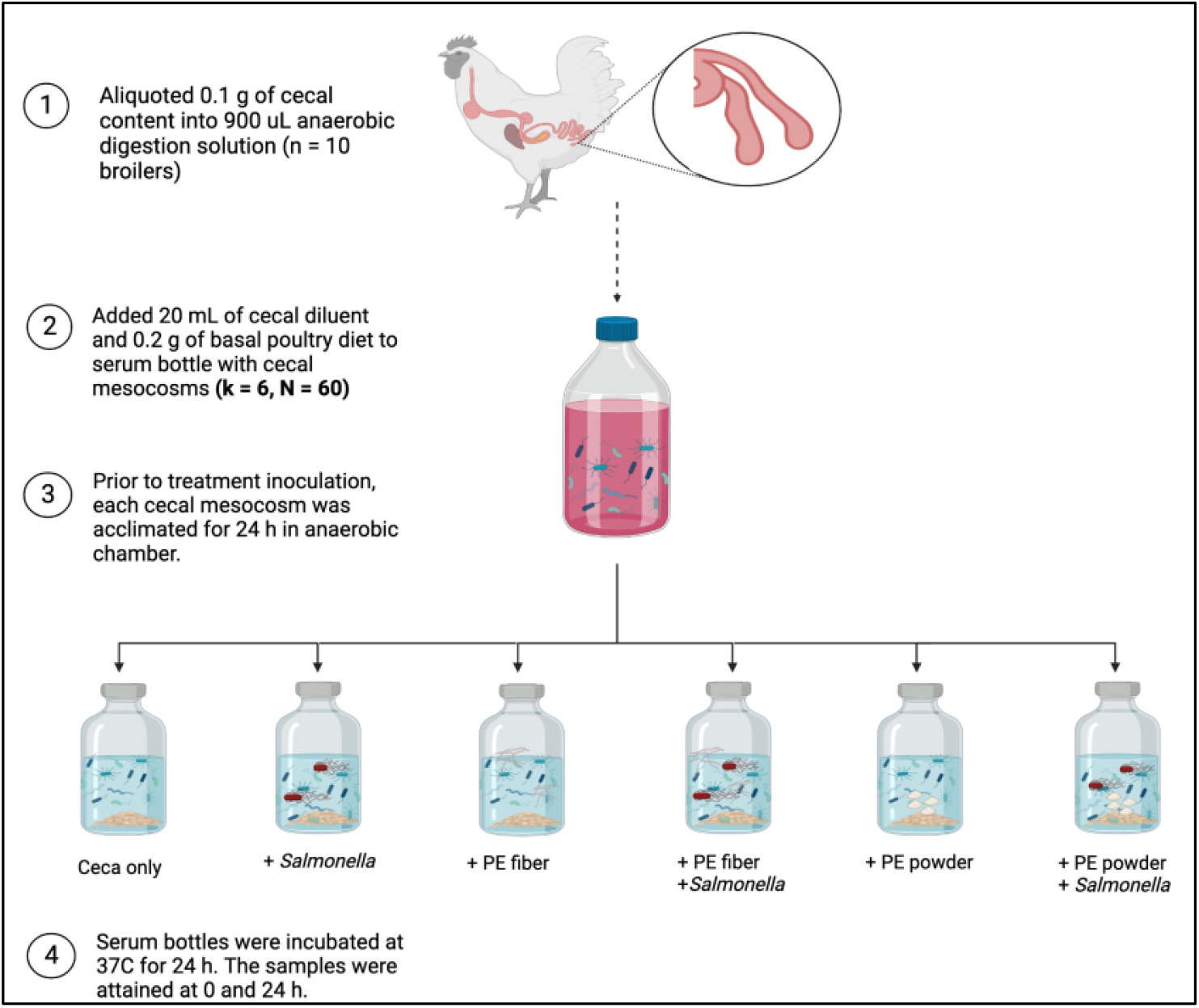
Design of anaerobic chicken cecal mesocosms.

Following the 24 h acclimation, designated polyethylene treatments were poured into the serum bottle containing cecal mesocosms. The *S*. Typhimurium was inoculated into the serum bottle containing cecal mesocosms using a pipette. The treatment groups were as follows: ceca only, *S*. Typhimurium only, polyethylene fiber (PE fiber), polyethylene fiber with *S*. Typhimurium (PE fiber + *Salmonella*), polyethene powder (PE powder) and polyethene powder with *S*. Typhimurium (PE powder + *Salmonella*). There were ten biological replicates per treatment group (six treatment groups; n= 10; N = 60). For the 0-hour time point, 1 mL of total contents was removed via serological pipette in duplicate and aliquoted into sterile microcentrifuge tubes, flash frozen in liquid nitrogen, and stored at −80°C. The serum bottle cecal mesocosms containing controls or treatments were incubated at 37°C for an exposure period of 24 h. At 24 hours, a second 1 mL sample was collected in duplicate and immediately stored in the same manner.

### Library Preparation for 16S rRNA Gene Amplicon Sequencing

Collected cecal mesocosm samples from each treatment group and each timepoint were stored in −80°C until genomic DNA extractions were performed. For gDNA extractions, 1 mL from each sample at both 0 h and 24 h timepoints were centrifuged for 5 min x 14000 rcf (N=120). The supernatant was then discarded, and DNA extraction was performed using the standard protocol for the DNeasy Blood and Tissue kit (Qiagen, Cat 69506). Total genomic DNA quality and concentration were verified using an Infinite 200Pro spectrophotometer (Tecan Nano Quant Plate^™^). The DNA extracts were diluted to 10 ng/µL in Buffer AE and stored at −80°C until further analysis.

To initiate amplicon library preparation for microbiome sequencing, DNA extracts were PCR-amplified with primers targeting the V4 region of the 16S rRNA gene. The primers were dual-indexed primers designed using high-fidelity polymerase, Pfx, according to the protocol by Kozich et al (33). Gel electrophoresis was performed to verify amplified PCR products. The amplified PCR products were then normalized to 20 µL using a SequalPrep Normalization kit (Life Technologies). Aliquots of 5 µL from each normalized sample were subsequently pooled into the final library. Final concentrations were verified using a KAPA library quantification kit (Kapa Biosystems) and a Qubit 2.0 Fluorometer (Invitrogen). Next, the final library was diluted to 20 nM with HT1 buffer and PhiX control v3 (20%, v/v) and 600 µL was loaded onto a MiSeq v2 (500 cycles) reagent cartridge (Illumina). The resulting sequences were uploaded to Illumina Sequence Hub and downloaded using BaseSpace Sequence Hub Downloader (Illumina).

### Microbiome Community Analyses

Quantitative Insights Into Microbial Ecology (QIIME) 2 (version 2022.2) (34) was utilized to perform microbiome bioinformatics. The raw Illumina amplicon sequence data was uploaded to BaseSapce Website (Illumina, San Diego, CA, United States) to assess sequence run quality and completion. The demultiplexed data was downloaded from the Illumina Basespace website and uploaded to QIIME2 using Casava 1.8 paired-end demultiplexed format (via QIIME import tools). Demultiplexed reads were then denoised with DADA2 (35) (via q2-dada2) and quality filtered using the chimera consensus pipeline. Next, the amplicon sequence variants (ASVs) were aligned using mafft (36) (via q2-alignment) and fastree2 (37) to produce the phylogenetic tree (via q2-phylogeny). Taxonomy was assigned to the ASVs in the feature table using (via q2-classifier sklearn) provided by QIIME2 with a confidence limit of 97% (38). The classifier was trained with SILVA 138 99% OTUs reference sequences (39). Alpha and Beta diversity analyses were then performed (via q2-diversity) for each time point.

Main effects and interactions were evaluated with ANOVA (via q2-longitudinal) (P-value ≤ 0.05;Q-value ≤ 0.05) (40) and ADONIS (R^2^ ≥ 0.50; P-value ≤ 0.05) (41). Pairwise comparisons using Kruskal-Wallis) was conducted for α-diversity metrics (Shannon’s Diversity index, Observed Features, Faith’s Phylogenetic Diversity, and Pielou’s Evenness) (P-value ≤ 0.05;Q-value ≤ 0.05) (42). In addition, pairwise comparisons using analysis of similarity analysis (ANOSIM) was performed for β-diversity metrics (Jaccard distance, Bray-Curtis distance, unweighted UniFrac distance and weighted UniFrac distance) (via q2-composition) (P-value ≤ 0.05;Q-value ≤ 0.05) (43, 44). Additional microbiome analyses (i.e., core microbiota and abundance-prevalence relationship) and data visualization (i.e., relative abundance bar plot and composition boxplots) in R (version 2022.12) were performed using an adapted protocol based on Sudakaran’s microbiome analysis tutorial (45). β-diversity metrics, Bray-Curtis and weighted UniFrac were visualized with principal component analysis plots in R.

### Metabolite Extraction

An aliquot of cecal mesocosm contents from each treatment and timepoint was used to determine protein concentration by a Bradford assay (46). Metabolites extraction was performed using 0.5 mL of from each cecal mesocosm sample (N = 105). Cells were lysed by three freeze-thaw and sonication cycles which consisted of thawing at room temperature for 10 min followed by a 30 second sonication on ice in a water bath sonicator (Branson 2800). A 1 mL aliquot of 2:2:1 (v/v) mixture of ≥ 99.9% purity acetonitrile (Fisher Scientific; Cat.A998-4) ≥ 99.8% purity methanol (Fisher Scientific; Cat. A412-4) and water was added to cell extracts and sonicated for 30 s and stored at −20°C overnight to allow cellular debris and protein to precipitate. The next day samples were centrifuged for 15 min at 20,784 x g at 4°C. The supernatant was transferred to new microcentrifuge tubes and dried for 4 h on a SpeedVac Concentrator (Thermo Scientific Savant DNA 120). Dried samples were then reconstituted in an acetonitrile: water (1:1 v/v) solvent mixture based on the normalized protein concentration of all samples, where the highest protein concentration sample had 100 µL of resuspension volume. Samples were vortexed for 30 s, sonicated on ice for 10 min and centrifuged for 15 min at 13,000 rpm at 4°C to remove any residual debris. The metabolite extracts were transferred into 0.3 mL HPLC autosampler vials with inserts (VWR International LLC, Cat. 9532S-1CP-RS) and stored in −80°C until analysis.

### Untargeted Metabolomics

Metabolite extracts were analyzed with ultra-high-performance liquid orbitrap chromatography mass spectrometry (UHPLC-MS) (Thermo Scientific Orbitrap Exploris 240 mass spectrometer). A Kinetex Core-Shell 100 Å column C-18 column (1 x 150 mm, 1.7 µm, Phenomenex) was used for separation of metabolites. Sample injection volume was 3 µL/min at a flow rate of 0.250 mL/min. Mobile phase A was composed of water with 0.1% formic acid, and mobile phase B was 0.1% in acetonitrile. The gradient began with 5% B from 0-3 min, then an increase and hold to 95% B until 18 min, followed by a decrease to 5% B at 18.50 min and hold at 5% B until 22 min. Data-dependent acquisition was used for the tandem MS workflow in positive ion mode.

### Untargeted Metabolomics Data Analysis

MetaboAnalyst 5.0 was used for statistical analysis and functional analysis of the MS1 data as described in Pang *et al.* (2022) (47). The raw files were converted to mzXML files and centroided (MS-1, orbitrap, positive mode) using ProteoWizard version 3.0 (48). Pairwise comparisons were performed to investigate potential differences of metabolite features between each treatment group at 0 h and 24 h and between the ceca only group and each treatment group. Then functional analysis (p-value = 1.0E-5, KEGG *E. Coli* library) was conducted to assess pathway-level changes to the cecal metabolome. Putative identification of metabolites was first carried out using MetaboAnalyst 5.0 which matches MS1 data compounds in Human Metabolome Database. Annotation and putative identification of metabolites was also performed on the raw MS/MS files in Compound Discoverer (version 3.3, Thermo Fisher Scientific) against mzCloud, mzVault, Metobolika and ChemSpider databases.

### Data Availability

Sequencing data were uploaded to NCBI’s Short Read Archive as BioProject ID PRJNA1043229 (http://www.ncbi.nlm.nih.gov/bioproject/1043229), and untargeted metabolomics data was uploaded to MetaboLights as project MTBLS9001 (www.ebi.ac.uk/metabolights/MTBLS9001).

## RESULTS

To determine the effects of plastic and pathogen co-exposure on the microbiome, ceca from 10 broiler chickens were harvested, diluted, aliquoted and grown overnight anaerobically. The 6 treatment groups were subsequently established containing ceca only (untreated), PE Fiber, PE powder, *S.* Typhimurium or a combination of each plastic with the pathogen. Cecal mesocosms were incubated for 24 hours. Samples were collected at 0 and 24 hours and analyzed for cecal microbial community composition and metabolome (Figure 1).

### Untreated cecal microbiome and metabolome were not significantly altered over 24-hour incubation

The ceca only group evaluated had the typical taxonomic composition and small molecule presence found within the ceca of broiler chickens. 16S rRNA gene amplicon sequencing targeting the V4 region was conducted, and the taxa were annotated to the genera level (Figure 2). The microbial composition detected in this study was found to be similar in membership to published 16S rRNA sequencing of cecal microbiomes from broiler chickens (49–51).

**Figure 2.**
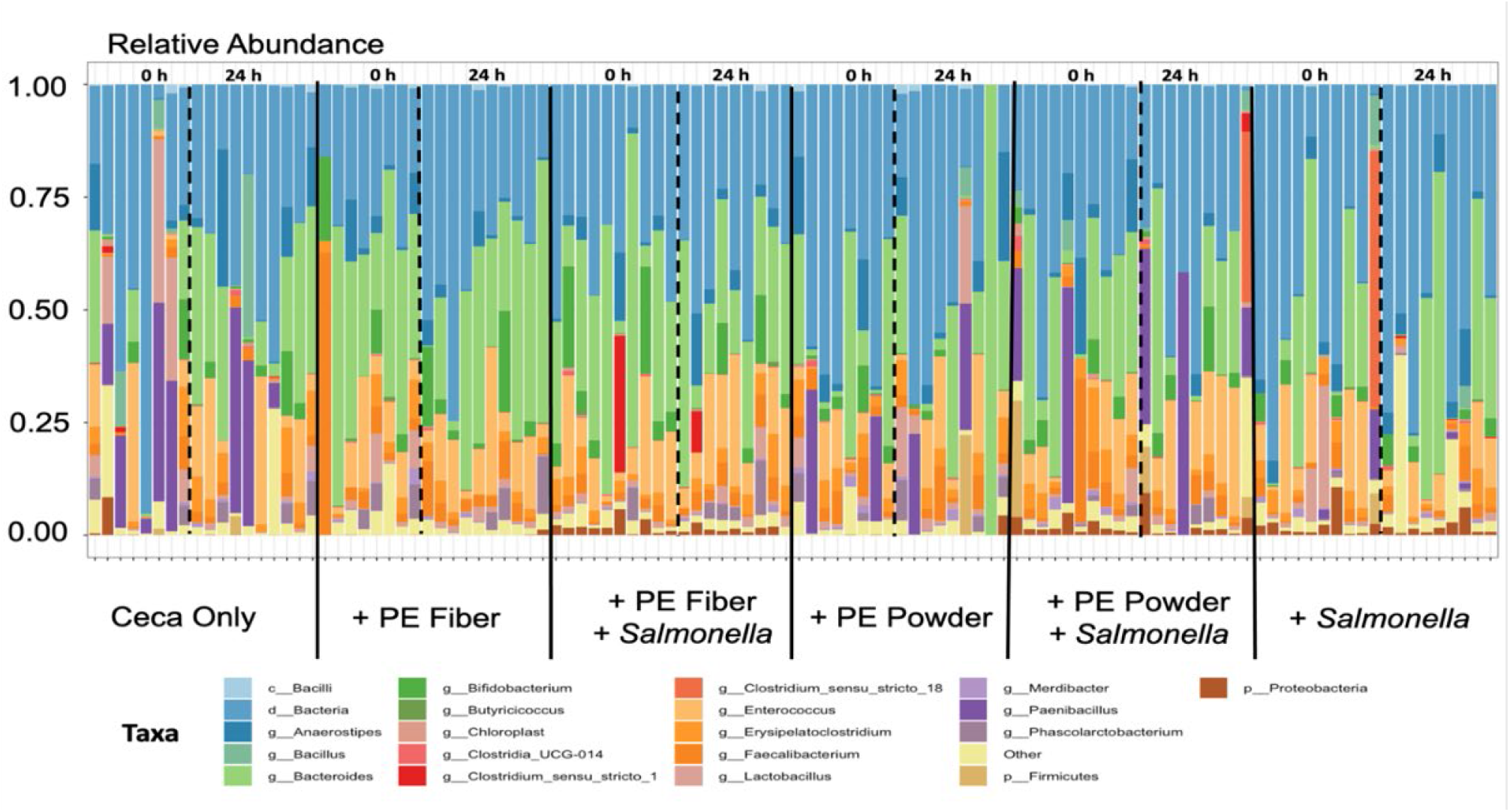
Taxa Bar Plot showing the Relative Abundance of the top 25 phyla by bird at 0 h and 24 h, separated by dashed line, for all treatment groups, separated by solid black line. The *Salmonella* added to the mesocosms is in the genus *Enterobacteria* and phylum *Proteobacteria* (brown bars).

To assess the effect of time on the cecal microbiome composition and metabolome, the untreated groups were compared at the 0- and 24-hour time points. The main effect “time” was significant for overall α-diversity metrics indicating phylogenetic diversity was altered over time (Table 1). However, when the 16S rRNA data was analyzed by time point (i.e., at 0 h and 24 h separately) there were no significant differences detected for α-diversity or β-diversity metrics for the untreated groups at the two timepoints (P-value ≥ 0.05) (Tables S4-S6). The relative abundance of the top 40 microbial taxa in individual samples was compared at the 0 h and 24 h timepoints (Figure 2) and found to have little variation with only four of 36 genera having a fold change (FC) in the relative of abundance greater than two: *Bacteroides, Enterococcus*, *Lactobacillus*, and an unclassified genus from the class Clostridia (Table S2). Similarly, untargeted metabolomics was used to assess the metabolome of the ceca. Pairwise comparison of metabolite features in ceca only samples at 0 h versus 24 h did not show any significantly dysregulated metabolites (P-value ≥ 0.05) among the 2,096 detected (Table S14). Overall, there were no significant changes in the cecal microbiome or metabolome of the ceca only group after 24 hours of incubation. This indicates that any observed changes in the treated conditions will be associated with the treatment and not normal fluctuation within the ceca.

**Table 1.** ANOVA results for alpha diversity metrics.

|  | Shannon | Pielou's Evenness | Faith* | Observed Features* |
| --- | --- | --- | --- | --- |
| PE | 0.071 | 0.057 | 0.717 | 0.717 |
| *Salmonella* | 0.701 | 0.823 | 0.449 | 0.449 |
| PE × *Salmonella* | 0.725 | 0.881 | 0.709 | 0.709 |
| Time | 0.243 | 0.798 | **0.042** | **0.042** |
| PE × Time | 0.800 | 0.782 | 0.974 | 0.974 |
| Time × *Salmonella* | 0.567 | 0.597 | 0.836 | 0.836 |
| PE × Time × *Salmonella* | 0.950 | 0.574 | 0.660 | 0.660 |
| *Bolded values indicate significant p-values. (N = 110; P-value ≤ 0.05) |  |  |  |  |

### *Salmonella enterica* Typhimurium influences the cecal metabolome

To assess the effect of *Salmonella* presence on chicken cecal microbiome and metabolome, groups containing *Salmonella* were compared to their respective no *Salmonella* group both at the 0-hour and then at the 24-hour timepoint (Ceca only vs + *Salmonella*; + PE Fiber vs + PE Fiber + *Salmonella*; and + PE Powder vs + PE Powder + *Salmonella*). For the microbial community composition, comparisons of α-diversity metrics were not found to be significant when analyzed at 0 h and 24 h (Kruskal-Wallis pairwise, Q-value ≥ 0.05) (Table S4a-d). Similar to α-diversity metrics, β-diversity metrics also showed insignificant results as determined by ANOSIM for each pairwise comparison (P-value ≥ 0.05; Q-value ≥ 0.05) (Table S5-6). Evaluation of the main effects and interactions as determined by ADONIS were also not significant (Table 7) (P-value ≥ 0.05; Q-value ≥ 0.05).

*Salmonella enterica* Typhimurium was inoculated into assigned cecal mesocosms to elucidate the effects of the enteric pathogen singularly and in conjunction with polyethylene. To verify the presence or absence of *S*. Typhimurium within inoculated cecal microbiome mesocosms, we analyzed the relative abundance of taxa for each treatment group. Comparing the relative abundance of bacterial phyla, we observed that the relative abundance of *Proteobacteria* is highest in *Salmonella*-containing groups (Figure 2, brown bars). As *Salmonella* is a member of the *Proteobacteria* phylum, this indicates that the pathogen inoculation was successful within our *Salmonella*-containing mesocosms. Furthermore, differential abundance testing using ANCOM at the genus level further confirmed this finding as *Enterobacterales* was the significant feature detected at 0 h between 24 h in *Salmonella*-containing groups (Table S3), indicating that *Salmonella* grew within the cecal mesocosms. To quantify the microbial composition changes over the time series, we analyzed the fold change in the relative abundance of all genera but evaluated the top 40 within the +*Salmonella* treatment group (Table S2). Within the +*Salmonella* group, the only two genera that exhibited fold change above two were *Enterococcus* and *Oscillibacter* with a fold change of 3.2 and 2.2, respectively (Table S2).

Although the microbial community composition was not significantly altered by the presence of *Salmonella* over a 24-hour period, activity of the microbiome, as measured by metabolome changes, was significantly altered. Global untargeted metabolomics was performed on all 6 treatment groups at both 0 h and 24 h timepoints. A multiple group comparison of the total metabolomes detected at 24 hours is plotted in Figure 2. Two clusters are observed which correspond to the presence or absence of *Salmonella*. This indicates that *Salmonella* substantially alters the metabolic activity of the cecal microbiome.

To determine metabolome similarities and dissimilarities between ceca only (untreated) and +*Salmonella* treatments, statistical analysis of the total metabolomes using pairwise comparisons was conducted (ceca only metabolome at 24 h vs +*Salmonella* metabolome at 24 h). We determined that 113 metabolites were significantly downregulated and 1 was significantly upregulated and 2,002 were not significantly dysregulated (Table S8). To ensure that significantly dysregulated metabolites were in fact correlated to *Salmonella* inoculation and activity, a comparison of the +*Salmonella* samples collected at 0 h and 24 h was analyzed. The result highlighted that 15 metabolites were significantly down-dysregulated, 139 metabolites significantly up-dysregulated and 1,971 were insignificant (Table S10). Significantly dysregulated metabolites associated with *Salmonella* inoculation putatively identifed with MetaboAnalyst included Simulanoquinoline, Asparaginyl-tryptophan, and Pyridoxamine (Figure 4a-c). Simulanoquinoline expression increased at 24 h for +PE fiber + *Salmonella*, +PE powder + *Salmonella* and +*Salmonella* groups (Figure 4a). In contrast, Asparaginyl-tryptophan expression decreased at 24 h within the +PE fiber + *Salmonella*, +PE powder + *Salmonella* and +*Salmonella* groups (Figure 4b). In addition, Pyridoxamine as identified by Compound Discoverer showed a maintained expression in *Salmonella*-containing groups (Figure 4c).

**Figure 3.**
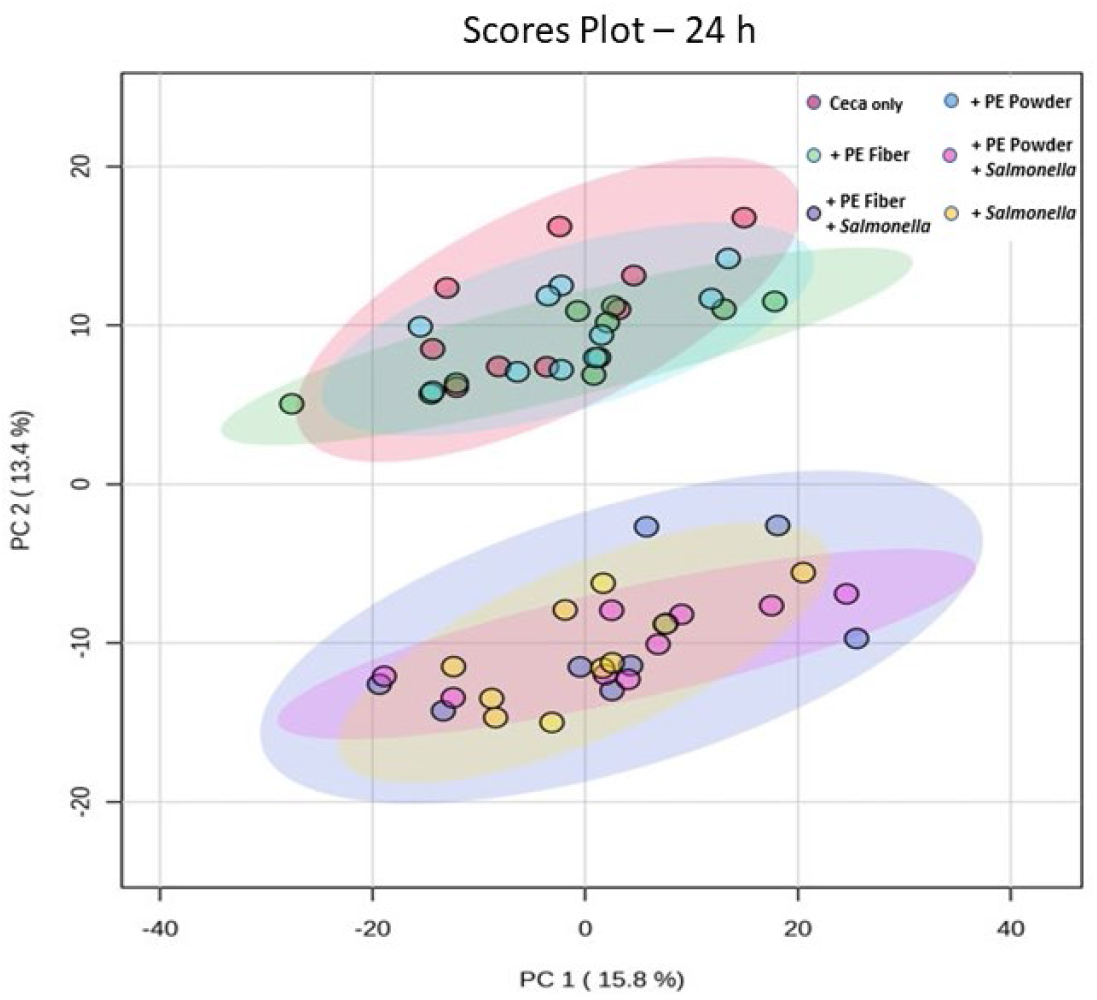
Principal component analysis of the metabolome at 24 h. Distinct clustering of treatment groups without *Salmonella* in upper portion of plot. The bottom portion shows a close relationship of *Salmonella*-containing treatment groups. This indicates that the presence of *Salmonella* has a greater influence on small molecule dysregulation than polyethylene fiber or polyethylene powder, singularly.

**Figure 4.**
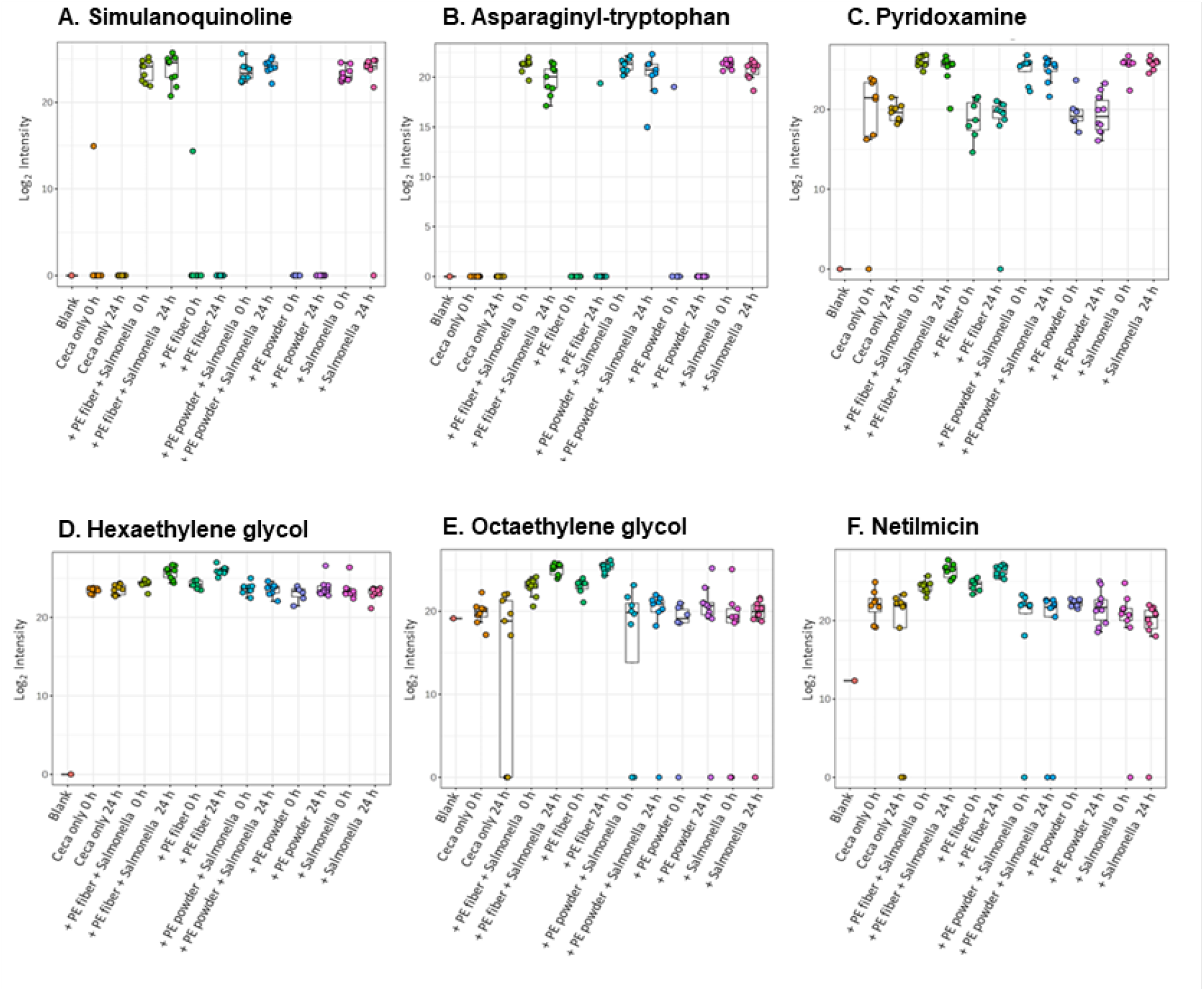
Significantly dysregulated metabolites associated with *Salmonella* inoculation or polyethylene fiber presence as determined by putative identification within MetaboAnalyst. Significantly dysregulated metabolites associated with the presence of *Salmonella* included A) Simulanoquinoline, B) Asparaginyl-tryptophan, and C) Pyridoxamine, Metabolites associated with PE fiber presence included D) Hexaethylene glycol, E) Octaethylene glycol and F) Netilmicin.

Following statistical analysis, we performed functional analysis to determine pathway-level changes that occurred within the metabolome. The Functional analysis module in MetaboAnalyst utilizes a modified Gene Set Enrichment Analysis and Mummichog algorithm (P-value = 1.0E-5, KEGG *E. coli* library) to map *m/z* features to metabolic pathways. The results demonstrated that the total metabolome of +*Salmonella* groups at 0 h and 24 h maintained a high expression of metabolites associated with arginine metabolism and porphyrin and chlorophyll metabolism but showed a decreased expression of biotin metabolism (213.1239_412.94 *m/z*_RT, where *m/z* is mass to charge ratio and RT is retention time in seconds for that metabolite feature) over the time course (Figure S4). There was also a noticeable increase in the feature 152.0347_371.77 which is associated with tryptophan metabolism (Figure S4). As expected, the ceca only metabolome at 24 h varied greatly from the +*Salmonella* metabolome at 24 h. For example, in the ceca only metabolome at 24h there was a low expression of features 259.1084_407.15 (porphyrin and chlorophyll metabolism), 155.0819_120.36 (Arginine metabolism), 213.1239_412.94 (biotin metabolism), 155.0818_91.01 (arginine metabolism) and 155.082_375.78 (arginine metabolism).

### The cecal microbiome is altered more by the PE fiber treatment than PE powder

To assess the effects of PE presence and polymer type on the cecal microbiome and metabolome, PE-containing groups were compared to the untreated ceca and the opposing PE treatment group (ceca only vs +PE powder; ceca only vs +PE fiber; +PE fiber vs +PE powder). Microbial community composition was analyzed by comparisons of α-diversity metrics using Kruskal-Wallis pairwise comparison in QIIME2. The results were not found to be significant for any treatment comparison at 0 h and 24 h (Q-value ≥ 0.05) (Table S4a-d). β-diversity metrics as measured by ANOSIM (P-value ≥ 0.05; Q-value ≥ 0.05) (Table S5-6) were also insignificant for each pairwise comparison. Evaluation of the main effects and interactions as determined by ADONIS were also not significant (Table S7) (P-value ≥ 0.05; Q-value ≥ 0.05). First, we assessed microbial composition variability among +PE fiber and +PE powder treatment groups. Interestingly, microbial composition of PE-containing groups as determined by calculating the mean relative abundance of phyla revealed that the composition of both +PE fiber and +PE powder consists of more *Firmicutes* than *Bacteroidetes* (Figure 5). Yet, at the genera level, the +PE fiber group had higher relative abundance of the genus *Bacteroides* (Figure 2).

**Figure 5.**
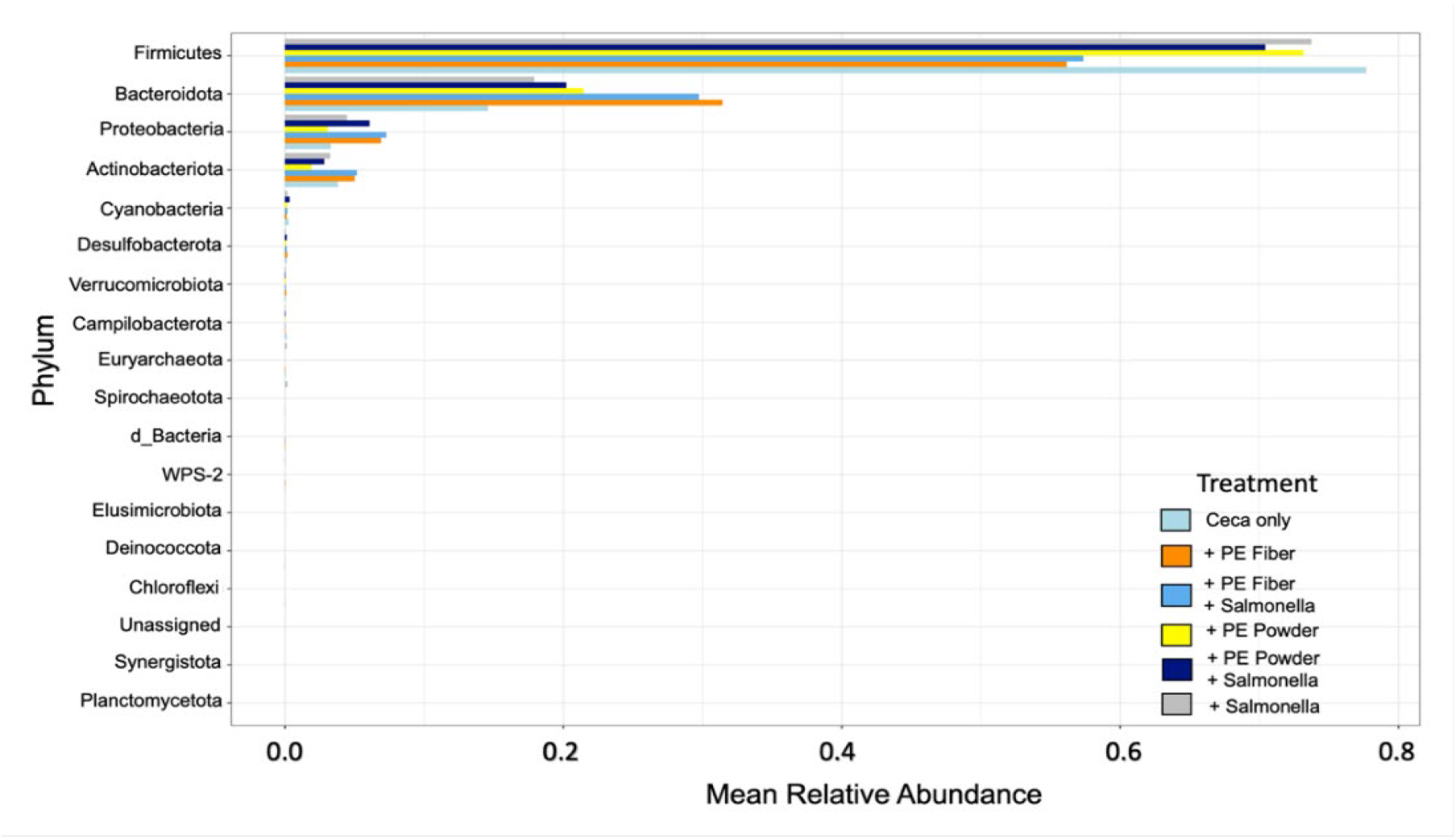
Mean relative abundance of phylum in each treatment group. Firmicutes, Bacteroidota, Proteobacteria, and Actinobacteria were the highest in mean relative abundance. Groups containing PE fiber have a lower mean relative abundance of Firmicutes and higher mean relative abundance of Bacteroidota compared to other treatment groups.

The effects of +PE powder on the cecal microbiome and metabolome were conducted by evaluating microbial composition over the time series. This was determined by calculating fold change in the relative abundance of genera for this treatment group. The top 40 genera were then analyzed. The largest fold change for the +PE powder group was for the genera *Bacteroides* and an uncultured bacterium (FC ≥ 2) (Table S2). The ceca only group also had a similar FC >2 for *Bacteroides* over the 24-hour time course. To determine the effects of PE powder on the total metabolome, a multigroup comparison was performed on the untargeted metabolomics data. Figure 3 highlights the similarities between +PE powder and +PE fiber as determined by principal component analysis. Further comparison of the ceca only metabolome at 24 h to the +PE powder total metabolome at 24 yielded no significantly dysregulated metabolites (P-value ≥ 0.05) (Table S14). To confirm this finding, comparison of +PE powder 0 h metabolome versus +PE powder 24 h metabolome was evaluated and also yielded no significantly dysregulated metabolites (P-value ≥ 0.05) (Table S14). Pathway-level changes, however; were comparable to +PE fiber and ceca only (Figure S4). In all, these results indicate the +PE powder treatment group did not significantly alter the cecal microbiome or metabolome.

To determine the effect of +PE fiber on the cecal microbiome and metabolome, +PE fiber containing samples were analyzed with the same workflow as +PE Powder groups. Despite insignificant results for microbial composition analysis, assessment of abundance level changes of genera indicated distinct changes in the microbial composition for the +PE fiber treatment groups. *Bifidobacterium* and *Sellimonas* increased at least 2-fold over the time course for the +PE fiber group, but not +PE powder (Table S2). Likewise, as mentioned above, +PE fiber groups had higher abundance of *Bacteroides* than other groups.

To assess the effects of PE fiber on the cecal metabolome, multigroup analysis and pairwise comparison (ceca only vs PE fiber; PE fiber at 0 h vs PE fiber at 24 h) was evaluated. Multiple group analysis of the total metabolome as determined with principal component analysis highlighted no distinct clustering of the total metabolome of +PE fiber at 24 h (Figure 3). Yet pairwise comparison of +PE fiber metabolome at 0 h to +PE fiber metabolome at 24 h yielded 49 significantly down-dysregulated metabolites, 4 significantly up-dysregulated and 1,998 insignificant metabolites (Table S9). Notably, Hexaethylene glycol, Octaethylene glycol and Netilmicin were metabolites associated with the presence of PE fiber as determined by putative identification within MetaboAnalyst (Figure 4). The same metabolites were significantly dysregulated for +PE fiber when comparing the ceca only metabolome at 24 h to the PE Fiber metabolome at 24 h (Table S9). Netilmicin, as annotated by Compound Discoverer, had an increased expression at 24 h for only +PE fiber and +PE fiber +*Salmonella* (Figure 4f). A similar trend was also detected for both hexaethylene glycol and octaethylene glycol (Figure 4d-e). This data suggests that PE fiber, but not powder leads to distinct changes in metabolic activity in the cecal microbiome.

As previously described, pathway-level changes were also assessed through utilization of the functional analysis module in MetaboAnalyst (P-value = 1.0E-5, KEGG *E. Coli* library). Comparison of the +PE fiber 0 h and 24 h metabolomes revealed that feature 259.1084_407.15 which is associated with porphyrin and chlorophyll metabolism was relatively lower within the total metabolome of +PE Fiber at 24 h (Figure S4). However, some individual +PE fiber samples show an increase in this feature. The +PE fiber metabolome at 24 h also had a low intensity of features 155.0819_120.36, 155.0818_91.01 and 155.082_375.78 which are associated with arginine metabolism (Figure S4). Surprisingly, comparison of ceca only at 24 h to +PE fiber at 24 h revealed similar metabolite levels among the treatment groups for the following features: 259.1084_407.15 (porphyrin and chlorophyll metabolism), 155.0819_120.36 (arginine metabolism), 155.0818_91.01 (arginine metabolism) and 155.082_375.78 (arginine metabolism) (Figure S4). Despite these similarities, the feature 165.0551_425.03 (phenylalanine metabolisms) showed higher intensity levels in +PE fiber at 24h when compared to ceca only metabolome at 24 h (Figure S4). These results indicate that the presence of PE fiber more than PE powder has an effect on the cecal metabolome and microbiome. However, unlike *Salmonella*-containing treatments, the presence of PE fiber affected the microbiome more than metabolome.

### Co-exposure to PE fiber & *Salmonella enterica* Typhimurium caused greater dysregulation in the cecal microbiome and metabolome

Groups containing both *Salmonella* and PE were compared to the untreated ceca and their respective no *Salmonella* group both at the 0 hour and 24-hour timepoint (Ceca only vs +PE fiber + *Salmonella*; Ceca only vs +PE powder + *Salmonella*; + PE Fiber vs + PE Fiber + *Salmonella*; and + PE Powder vs + PE Powder + *Salmonella*) to assess the effect of both PE and *Salmonella* presence on chicken cecal microbiome and metabolome. Microbial composition analysis as determined by Kruskal-Wallis pairwise analysis of α-diversity metrics was not significant at 0 h and 24 h for any comparison (Q-value ≥ 0.5) (Table S4a-d). β-diversity metrics were evaluated with at 0 h and 24 h, and the results were also insignificant for each pairwise comparison (R^2^ ≥ 0.5; Q-value ≥ 0.5) (Table S5-6). In addition, overall assessment of dissimilarities among the treatment groups as determined by Bray-Curtis and Weighted UniFrac PCoA plots did not demonstrate any distinct clustering (Figure S1). Although α-diversity and β-diversity metrics were insignificant, overall microbial composition did vary slightly among co-exposed +PE fiber + *Salmonella* and +PE powder + *Salmonella* treatment groups compared to individual exposure treatments. For instance, there was a lower relative abundance of *Paenibacillus* in the +PE fiber + *Salmonella* but not in the +PE powder + *Salmonella* group (Figure S2).

To quantify abundance level changes in the cecal microbiome resulting from +PE fiber +*Salmonella* exposure, we calculated the fold change (FC) in relative abundance of genera. FC in the relative abundance of the top 40 genera revealed that *Lactobacillus* was the only genera within the +PE fiber +*Salmonella* group which increased greatly (FC >2) (Table S2). Remarkably, the only other treatment groups that had a FC > 2 for *Lactobacillus* was the ceca only treatment group (Table S1). In addition, the *Firmicutes/Bacteroidetes* ratio (F/B ratio) was evaluated to determine if the +PE fiber + *Salmonella* treatment led to dysbiosis of the cecal microbiome. Table 2 highlights that the presence +PE fiber + *Salmonella* results in an F*/B* ratio of 1.93. Conversely, the F/B ratio was 5.30 for the ceca only group, which indicates that +PE fiber +*Salmonella* exposure resulted in greater disruption to the cecal microbiome (Table 2).

**Table 2.**
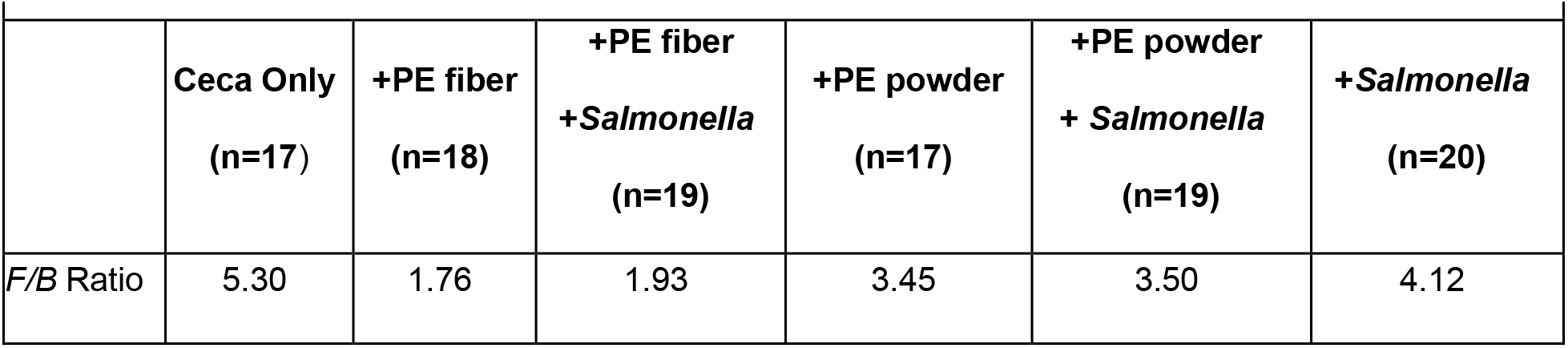
*Firmicutes/Bacteroides* Ratio for each treatment group overall.

|  | Ceca Only<br>(n=17) | +PE fiber<br>(n=18) | +PE fiber<br>+*Salmonella*<br>(n=19) | +PE powder<br>(n=17) | +PE powder<br>+ *Salmonella*<br>(n=19) | +*Salmonella*<br>(n=20) |
| --- | --- | --- | --- | --- | --- | --- |
| F/B Ratio | 5.30 | 1.76 | 1.93 | 3.45 | 3.50 | 4.12 |

Next, we evaluated the presence of +PE fiber +*Salmonella* on the cecal metabolome using multiple group analysis and pairwise comparison (ceca only vs +PE fiber +*Salmonella*; +PE fiber +*Salmonella* at 0 h vs +PE fiber +*Salmonella* at 24 h). Activity of the microbiome, as measured by metabolome changes, was significantly altered by the presence of +PE fiber +*Salmonella* but not +PE powder +*Salmonella.* As mentioned previously, multiple group analysis as determined principal component analysis showed distinct clustering of *Salmonella*-containing groups (Figure 3). Pairwise comparison of the ceca only vs +PE fiber +*Salmonella* at 0 h and 24 h revealed significantly dysregulated metabolites associated with the presence of both PE fiber and *Salmonella.* Pairwise comparison of ceca only at 24 h to +PE fiber +*Salmonella* metabolome at 24 h yielded 131 significantly down-dysregulated metabolites, 9 significantly up-dysregulated metabolites and 1,973 insignificant metabolites (Table S12). In addition, a pairwise comparison of +PE fiber +*Salmonella* at 0 h vs +PE fiber +*Salmonella* at 24 h yielded 19 significantly down-dysregulated metabolites, 130 significantly up-dysregulated metabolites and 1,976 insignificant metabolites (Table S12).

Functional analysis was then performed to assess pathway-level changes that occurred within the metabolome of +PE fiber +*Salmonella* (P-value = 1.0E-5, KEGG *E. Coli* library). Pathway analysis results for +PE fiber + *Salmonella* treatment groups were comparable to the +*Salmonella* group. For instance, there was also a decrease in feature 213.1239_412.94 at 24 h which is associated with biotin metabolism (Figure S4). Furthermore, there was a consistently higher level of metabolites 155.0819_120.36 (arginine metabolism), 155.0818_91.01 (arginine metabolism), 155.082_375.78 (arginine metabolism), and 259.1084_407.15 (porphyrin metabolism) (Figure S4).

The same analyses were performed as above to assess overall microbial composition and abundance level changes in the +PE powder +*Salmonella* treatment group. Unlike the +PE fiber +*Salmonella* treatment group, *Enterococcus, Phascolarctobacterium*, *Lactobacillus*, and two unassigned bacteria had a FC >2 (Table S2). Additionally, +PE powder +*Salmonella* had F*/B* ratio of 3.50 which is lower than ceca only and +*Salmonella* treatment groups but higher than +PE fiber +*Salmonella* (Table 2). Therefore, demonstrating that +PE powder +*Salmonella* results in minimal disruption of the cecal microbiome compared to its counterpart, +PE fiber +*Salmonella*.

To assess the effects of +PE powder +*Salmonella* on the cecal metabolome, multigroup analysis and pairwise comparison (ceca only vs +PE powder +*Salmonella*; +PE powder +*Salmonella* at 0 h vs +PE powder +*Salmonella* at 24 h; +PE fiber +*Salmonella* at 0 h vs +PE fiber +*Salmonella* at 24 h) was evaluated. Again, multiple group analysis as determined principal component analysis showed distinct clustering of +PE powder +*Salmonella,* +PE fiber +*Salmonella* and +*Salmonella* (Figure 3). Pairwise comparison of ceca only to co-exposed +PE powder +*Salmonella* at 24 h detected 112 significantly down-dysregulated metabolites, 1 significantly up-dysregulated metabolite and 1,993 insignificant metabolites (Table S13). While pairwise comparison of the +PE powder +*Salmonella* metabolomes at 0 h vs 24 h resulted in 14 significantly down-dysregulated metabolites, 148 significantly up-dysregulated metabolites and 1,963 insignificant metabolites (Table S13). Following statistical analysis, we performed functional analysis to more accurately determine pathway-level changes that occurred within the +PE powder +*Salmonella* metabolome (P-value = 1.0E-5, KEGG *E. Coli* library). Metabolite levels were similar to the +PE fiber +*Salmonella* and +*Salmonella* total metabolomes where features such as 155.0819_120.36 (arginine metabolism), and 259.1084_407.15 (porphyrin metabolism) had persistent high levels over the time series (Figure S4). Due to the lack of microbial activity changes detected in the +PE powder metabolome, pathway-level changes observed in the +PE powder +*Salmonella* metabolome are likely due to *Salmonella* causing greater disruption in the cecal microbiome and metabolome.

## DISCUSSION

This study focused on determining the effects of a 24 h co-exposure of low-density PE MPs and *S.* Typhimurium. Several analyses were conducted to establish if the main effects of “time,” “polyethylene” and “*Salmonella*” resulted in significant disruption to taxonomic composition and metabolic activity within the cecal microbiome. The results indicated that for α-diversity and β-diversity metrics there was no significant effect on community composition, richness, or diversity by the main effects: “time,” “polyethylene” and “*Salmonella*.”

Although α-diversity and β-diversity metrics were insignificant for the community overall, there were treatment specific responses in the relative abundance of a few taxa. In the +*Salmonella* treatment group, the only genera that responded were *Enterococcus* and *Oscillibacter* (Table S2). Previous research by Khan and Chousalkar (2020) reported that *Salmonella* infection in chickens resulted in a significant increase in abundance for several genera including *Oscillibacter,* which resulted in the birds experiencing gut dysbiosis (52). Our results are consistent with this finding and suggest that *Salmonella* inoculation may lead to dysbiosis.

+PE fiber induced changes to the cecal microbiome were in *Bifidobacterium* and *Sellimonas* (Table S2). *Bifidobacterium* has been reported to be increased in abundance following oral ingestion of the dietary fibe*r, Inulin,* which prevented *Salmonella* colonization in chickens (53). Interestingly, in our findings, plastic fiber exposure also increased the presence of beneficial bacteria even though we assume that PE fiber is non-fermentable by cecal microorganisms. This suggests that fiber presence, regardless of material, stimulates beneficial microbial activity. In the co-exposure treatment with PE fiber and *Salmonella*, both *Oscillibacter* and *Bifidobacterium* had decreased abundance over the 24 hours. This suggests that there are interactions between the pathogen and the microplastic and that these interactions are changing how the exposures affect the cecal microbial community. In this case, both beneficial and harmful species were inhibited in growth, highlighting the complexity of the multifaceted interactions between contaminants and gut microbial communities. The health outcomes from this co-exposure then depend on which factor induces stronger changes.

+PE powder did not elicit any significant responses. However, in this study, the ceca only and +PE powder groups were the only treatments with a FC > 2 for *Bacteroides*. Jacobson *et. al* (2018) reported that propionate produced by *Bacteroides* prevented *Salmonella* colonization in mice (60). Assuming propionate is being produced by the *Bacteroides* within the +PE powder group, this would explain the less drastic effect of PE powder on cecal taxonomic composition. In addition, the surface texture of PE powder may not allow for microbial colonization which would prevent changes to cecal microbial composition (60). This is an indication that polymer type, surface morphology and size affect the growth and abundance of microbial communities. As a result, intestinal homeostasis was affected by +PE fiber and +PE fiber + *Salmonella* treatments, but not +PE powder and +PE powder + *Salmonella* treatments. Therefore, despite insignificant microbiome community analyses, microorganisms were greatly altered by the presence of +PE fiber and +PE fiber *+ Salmonella,* but not +PE power and +PE power + *Salmonella*.

In addition to the response of specific genera to treatment, the *Firmicutes/Bacteroides* ratio is an important metric commonly used with other health markers to determine the presence of gut dysbiosis or gastrointestinal disorders. Specifically, an increased *Firmicutes/Bacteroides* ratios is correlated to obesity while a decreased *Firmicutes/Bacteroides* ratio is associated with Irritable Bowel Disease and other chronic inflammatory disorders (54, 55). This trend has been reported in several studies profiling the microbiota of Chron’s and colitis samples (54, 56–58). In contrast, there are studies which have reported no inverse relationship of *Firmicutes* and *Bacteroides* (55, 59). Despite this contradiction, the *Firmicutes/Bacteroides* ratio has long served as an acceptable way to assess gut health as these are the dominant bacterial phyla in many species including chickens (45, 50). In this study, *Firmicutes*, *Bacteroidota* and *Actinobacteriota* were confirmed as the prevalent phyla (Figure S3). The *Firmicutes/Bacteroides* ratio for each treatment group was calculated and revealed that the mean relative abundance of phyla was greatly affected by the presence of PE fiber with and without *Salmonella* (Table 2). +PE fiber and co-exposure +PE fiber + *Salmonella* treatment groups had *Firmicutes/Bacteroides* ratios much lower than ceca only, +PE powder, +PE powder +*Salmonella* and +*Salmonella*. The lower *Firmicutes/Bacteroides* ratio in PE fiber containing groups was from a decrease in *Firmicutes* and an increase in *Bacteroides* over the 24-hour exposure. Similar to our observations, Sun et al. (2021) also reported a decrease in *Firmicutes* and increase in *Bacteroides* in the gut microbiome of mice following oral exposure to PE microplastics, although the plastic shape was not reported (4). We further speculate that the continued growth of *Firmicutes* within the +PE powder and +PE powder + *Salmonella* groups (higher *Firmicutes/Bacteroides* ratios) but not the +PE fiber and +PE fiber + *Salmonella* groups is a result of increased interference to microbial interactions both synergistic, and antagonistic by +PE fiber + *Salmonella*. Therefore, our findings and those from studies in mammals indicate that the presence of PE fiber and *Salmonella* may result in animal gut dysbiosis. These observations also highlight the complexity of determining exposure effects because of the large number of factors and their interactions. This suggests that studies of individual exposures or studies conducted in highly controlled settings may not reflect the actual health outcomes from environmental exposures.

Untargeted metabolomics analyses used to determine metabolic activity in the cecal microbiome revealed *Salmonella* had a greater influence on the cecal metabolome than other treatments. Furthermore, assessment of metabolic activity was an important factor in understanding the interaction of PE fiber and PE powder with and without *S.* Typhimurium. Distinct metabolites associated with the presence of +PE fiber and +*Salmonella* treatments were observed. Significantly dysregulated compounds associated with *Salmonella* presence included simulanoquinoline, asparaginyl-tryptophan and pyridoxamine (Figure 3a-c). The first, simulanoquinoline, has been reported as a cytochrome P450 3A4 (CYP3A4) inhibitor and an α-glucosidase inhibitor (38, 39). CYP3A4 has been detected in human and chicken gastrointestinal tracts and livers and is known to be responsible for phase 1 metabolism of xenobiotics, bile acids, antibiotics, and dietary compounds (38, 40). Within this study, simulanoquinoline levels increased over the time course for +PE fiber +*Salmonella*, +PE powder +*Salmonella* and +*Salmonella* treatment groups (Figure 3a), suggesting that *Salmonella* produced this metabolite, stimulated production of it or induced the release of the molecule from the feed particles that were in the mesocosm. Daou et al (2022) reported detecting simulanoquinoline in abundance in the crude extracts of the shrub *Tamarix nilotica* (61). Given that the basal diet incorporated into the cecal mesocosms contains plant compounds, it is plausible that interactions among the PE fiber, *S*. Typhimurium, and ingredients such as soybean meal in the basal diet results in generation of this metabolite.

Asparaginyl-tryptophan was another metabolite that was significantly dysregulated when *Salmonella* was present. Genes associated with tryptophan biosynthesis and transport have been reported to be essential for *Salmonella* biofilm formation and attachment (62). *Salmonella enterica* Typhimurium, specifically, has been documented to attach and thrive on abiotic and biotic surfaces (26, 63, 64). As such, the upregulation of asparaginyl-tryptophan in *Salmonella*-containing groups is likely a result of *S.* Typhimurium biofilm formation and attachment to either PE fiber, PE powder or the serum bottles used for the cecal mesocosms. Unfortunately, we were unable to recover the PE fiber or powder from the cecal mesocosm to confirm presence or absence of biofilms. We did perform a separate *in vitro* experiment where we grew *S*. Typhimurium without the cecal community in the presence of the same PE fiber and PE powder and stained for biofilm formation (Figure S6). This quantitative biofilm formation assay revealed that *Salmonella* formed more biofilms on our 50 µm PE fibers compared to PE powder, consistent with literature reports of Salmonella forming biofilms on different plastic surfaces (63, 64). Considering the increased relative abundance of *S.* Typhimurium in the presence of PE fiber, the significantly dysregulated metabolites associated with *Salmonella* and the higher biofilm presence on PE fiber microplastics, it suggests that PE fiber presence and *S.* Typhimurium biofilm formation potential are essential factors mediating the effects of this co-exposure. In addition, biofilm formation on PE MPs may explain why the co-exposure had a greater disruption, as seen with the mean relative abundance of phyla, than the simple sum of the individual exposures (Figure 2).

Pyridoxamine, a form of vitamin B6 (65) was detected in this study at the highest levels in *Salmonella*-containing groups: +PE fiber + *Salmonella,* +PE powder + *Salmonella* and + *Salmonella* (Figure 3c). However, the concentration of pyridoxamine was maintained indicating an accumulation of this compound within these groups. The presence of this compound is likely due to the premix added in the poultry basal diet which contained 4 mg/kg of vitamin B6 (Table S7). Pyridoxamine accumulation within *Salmonella-*containing groups is likely a result of *S.* Typhimurium lacking a periplasmic binding protein essential for vitamin B6 synthesis (58, 65). Therefore, higher concentrations of pyridoxamine are detected in *Salmonella*-containing treatment groups because *Salmonella* cannot consume it whereas without the presence of *Salmonella* other members of the cecal community appear to metabolize it. In all, each of the metabolites highlighted were upregulated in the presence of *S.* Typhimurium, thus, providing evidence that *Salmonella* greatly influences the cecal metabolome.

In addition to metabolites linked to *Salmonella*, there were significantly dysregulated metabolites associated with +PE fiber presence, namely hexaethylene glycol and octaethylene glycol (Figure 3d-3f). These molecules are a part of a class of organic compounds called polyethylene glycols (65). Hexaethylene glycol and octaethylene glycol were also significantly down-regulated in a pairwise comparison of the +PE fiber samples collected at 0 h and 24 h (Table S9). The same metabolites were also significantly up-regulated in pairwise comparison of +PE fiber at 24 h to +PE powder at 24h (Table S9). This indicates that the metabolites were not present in high quantities at the initial time of exposure and may be the result of chemicals or other unknown compounds leaching from PE fiber. Since our exposure was commercial plastics and not pure polyethene polymer, water soluble additives and plasticizers in the plastic will leach into the cecal matrix. While these molecules could be at least partially metabolized by cecal microbiome members, we did not detect degradation products of these compounds over the 24-hour exposure, likely because the microbiome is replete with preferential nutrients from the poultry basal diet.

Netilmicin was also upregulated in the +PE fiber treatment group. Netilmicin is a compound known to have bactericidal effects in vulnerable organisms (65). We suspect that netilmicin originated from the commercial poultry basal diet that was added into the cecal mesocosm as a nutrient source during the acclimation period, however, the feed was not characterized prior to its use in the cecal mesocosms. It may also have originated from the cecal microbial community during the acclimation phase as antibiotic production is part of typical microbial interactions in a complex community. The metabolite is highest in the presence of PE fiber with and without *Salmonella* but not PE powder with and without *Salmonella*. This could suggest that the presence of PE fiber induced microbial interactions that led to the production of this antibiotic. This further indicates that treatment type influences how microbes within the cecal mesocosms utilize metabolites, especially those derived from the commercial poultry basal diet.

In all, statistical analyses of the total metabolome of each treatment group highlight that greater dysregulation occurs in the presence of co-exposure to +PE fiber + *Salmonella* (Table S7-13). To better assess this co-exposure future evaluation should incorporate a longer exposure duration, analysis of *Salmonella* biofilm formation pre- and post-inoculation and characterization of nutritional sources such as the poultry basal diet utilized in this study. Regardless of these limitations, the results of this study provided evidence that co-exposure to +PE fiber and *S.* Typhimurium but not PE powder, leads to greater dysregulation of the cecal microbiome and metabolome of broiler chickens.

## CONCLUSION

By integrating 16S rRNA gene amplicon microbial community sequencing and untargeted metabolomics, we investigated the response of the ceca microbiota and its related metabolites following exposure to low-density PE and *S.* Typhimurium in mesocosms. PE in the fiber but not powder form caused disruption to microbial community membership with both positive and negative indicators of gut health. The presence of *Salmonella* altered the metabolome with very few changes in microbial community member relative abundance suggesting that *Salmonella* excretes unique metabolites or that the cecal community metabolism is responding to *Salmonella* inoculation. We determined that the co-exposure of +PE fiber + *Salmonella* yielded greater disruption to the cecal metabolome and microbiome than either individual treatment. This indicates that polymer shape and size affect the growth, metabolism, and abundance of microbial communities in chicken ceca. Additionally, effects of individual exposures were contaminant specific. In all, co-exposure to PE fiber and *S.* Typhimurium appears to modulate the cecal microbiome through MPs-microbial interactions which led to microbial composition imbalance and altered metabolic activity.

## Acknowledgements

The authors gratefully acknowledge use of facilities and instrumentation at the UW-Madison Wisconsin Centers for Nanoscale Technology (wcnt.wisc.edu) partially supported by the NSF through the University of Wisconsin Materials Research Science and Engineering Center (DMR-1720415). We would also like to thank Dr. Bradley Bolling and Klay Liu for providing access to the mass spectrometer and the helpful discussions on untargeted metabolomics. In addition, we thank the University of Wisconsin Madison’s Poultry Research Facility and staff members for managing the husbandry of the broilers acquired for this study. The author CC would like to acknowledge the University of Wisconsin-Madison’s Molecular and Environmental Toxicology graduate program for its financial support through the Molecular and Environmental Toxicology Training Award (T32 ES007015) as well as Fuad Shatara for preparing the polyethylene fiber utilized in this study. The author EGO would like to acknowledge the USDA Hatch Grant (project # AAI9572; award: MSN248836) for its financial support.

